# Whole genome sequencing of hematologically stained cells catapulted from Cell smears

**DOI:** 10.1101/2021.12.31.474675

**Authors:** Sangwook Bae, Yushin Jung, Sungsik Kim, Jinhyun Kim, Amos Chungwon Lee, Dongsoon Lee, Sunghoon Kwon

## Abstract

Analyzing archived peripheral blood smears is a potential route towards gaining cell morphology and genome information of blood cell types from various diseases. Yet, acquiring whole genome information from morphologically targeted cells was difficult, especially for rare cell types. The main causes for such difficulty were the inevitable usage of cell stains leading to whole genome amplification inhibition, and insufficient cell isolation performance of previously introduced laser microdissection (LMD) techniques. Here, we introduce a new laser-based cell isolation technique and a whole genome amplification (WGA) protocol optimized for whole genome analysis from minute input of hematologically stained cells. We were able to perform whole genome copy number profiling and SNP analysis from as little as 5 cells.

## Introduction

Peripheral blood smears provide cell morphology-based information at the single cell level. These cytology specimens are used for counting morphologically abnormal cells, or measuring the composition of heterogeneous cell population^1, 2^. When diagnosing leukemia or lymphoma, these samples guide additional tests needed and treatment options, thus reducing potential waste of resources. And since cell smears well preserve cell morphology and genomic contents over long periods, patient cell smears are often archived in large hospitals. Therefore, these archives are potential route to large scale genomic studies.

Many genetic tests analyze mutations in only a small number of genes known to be related to disease of interest^3^. But such gene set selection becomes difficult when there exists patient-to-patient heterogeneity or variance in mode of disease inheritance^4–6^. Although many genetic tests have been introduced such as FISH^3,7–9^ and in-situ sequencing^10–12^ adapting whole exome sequencing (WES) or even whole genome sequencing (WGS) is becoming the most desirable approach because they provide both high sensitivity and coverage^13^. Furthermore, when investigating cancer and other diseases that may have occurred from cells with rare genomic instabilities (e.g. somatic mutation), it becomes important to enable WGS even with small number of cells. Therefore, future healthcare systems will greatly benefit from low-cell-input WGS applied to archived blood smear cells.

Performing genetic analysis on smeared cells usually requires transferring target cells on slide glass into small volume suspensions, which is usually done with infrared (IR) laser-based microdissection (LMD). But LMD systems, whether UV or IR mode, requires contour dissection that frequently leads to target cell damage or neighboring cell contamination when applying to compact tissues and incompletely dissociated cytological specimens^14–17^. And due to other technical hurdles, previous genetic analysis on such LMD-isolated blood smear cells were mostly multiplexed PCR and sequencing^14, 18,19^ and, to our knowledge, WGS reports are still lacking. There are two hurdles we need to overcome. First, cell stains, such as H&E, which are indispensable for morphology analysis are known to hamper DNA amplification efficiency^20–23^ unless amplification is preceded by DNA purification^24–25^. But most DNA purification methods are not feasible for minute cell input. So a robust whole genome amplification (WGA) protocol that produces whole genome coverage even from a small number of stained cells is needed. Second, when isolating cells from glass slides, IR LMD suffers from low cell isolation resolution (7.5μm laser diameter^14^) compared to UV based LMD, which could lead to retrieval of off-target cells when applying to compact tissues, or incompletely dissociated cytological specimens^15, 16^. Previously, our group introduced a high throughput IR laser-aided cell isolation and sequencing system that obtained higher cell isolation throughput and better genomic DNA quality than conventional LMD or laser pressure catapulting (LPC) systems^26–27^.

Based on our previous invention, here, we introduce a dissection-free, single cell laser isolation system and a WGA protocol optimized for blood smear samples. The isolation system increases the isolating speed and resolution of the single cell-catapulting force while minimizing damage of laser-isolated cells. We introduce a guideline of cell isolation, cell lysis, and WGA that allows whole genome copy number (CN) profiling and SNP detection from samples containing as little as 5 blood smear cells. We provide information regarding which cell staining material is preferred, the minimum number of cells required, and proper blood smear storage condition for high quality whole genome analysis similar to bulk whole genome sequencing. We remark that the entire workflow is adaptable to routine blood smear diagnosis.

## Materials and Methods

### Materials

ITO coated glass was ordered from Fine Chemical Industry, with different sputtering thickness of ITO coating layer; 150 nm and 300 nm. Trypan blue (Cat. no. T8154), Accustain Giemsa (Cat. no. GS500) and Wright Giemsa (Cat. no. WG16) was purchased from Sigma Aldrich. Hemacolor was purchased from Merck (Cat. no. 111661). Whole genome amplification kit that contains sample buffer, denaturation solution, neutralization buffer, reaction buffer, and MDA enzyme mix was purchased from GE (Illustra Genomiphi V2 DNA amplification kits, Cat. no. 25-6600-30). SYBR green I was purchased from Life Technologies (Cat. no. S7563). PCR master mix was purchased from the New England Biolabs (Quick-Load® *Taq* 2X Master Mix) or from KAPA Biosystems (KAPA HiFi HotStart ReadyMix, 2X). DNA purification kit was purchased from Beckman Coulter (Agencourt AMPure XP kit, Cat. no. A63880). Proteinase K was purchased from Sigma Aldrich (Cat. no. P4850-1ML).

### Cell samples

Blood samples and FISH stained samples were provided by prof. Lee at SNUH. Tissue stamped sample was provided by prof. Han at SNUH. HL60 cell line was cultured as recommended in the ATCC standard protocol. In brief, cells were cultured in Iscove’s Modified Dulbecco’s Medium (IMDM) supplemented with 20% FBS and 1% Penicillin-Streptomycin. Cells were cultured in a 37°C, 5% CO_2_ conditioned incubator. Cell medium was changed to fresh medium every 2~3 days to maintain cell concentration of 10^5^~10^6^ cells/mL.

### Blood smear

A 10uL drop of blood or cell sample (10^5^-10^6^ cells / mL) was dispensed near the corner of a bare glass slide or an ITO glass slide. Another glass slide was held on a 45° angle towards the sample droplet. While maintaining contact with the bottom slide, the top glass was pulled back to contact the drop. Immediately after droplet contact, the top glass slide was pushed forward in one motion while maintain firm contact and angle with the bottom slide. The resulting smear was left at ambient condition until fully dried.

### Cell staining

We performed two different Giemsa staining methods; one acquired from SNUH and the other from Sigma-Aldrich. We also performed Hemacolor staining and Trypan Blue staining method as well. Staining, other than the Giemsa from SNUH, was performed as suggested by the manufacturer. Giemsa staining protocol from SNUH was as follows. Cells were first smeared on bare glass or ITO glass using the blood smear method described above. After air drying, the slide was dipped into 100 % methanol for 30 seconds for fixation. Immediately, the slide was then dipped in Giemsa staining solution for 7 minutes. Then it was rinsed with phosphate-buffer saline (PBS) solution and then with deionized water. The slide was then fully dried in ambient condition and stored in −20°C until further experiment.

### Laser microdissection and lysis of cells

Pulse width of the IR laser was 7~10 nanoseconds with wavelength 1064 nm. For the bare glass experiment, we first ablated the surrounding contour of the target cell and isolated the cell by a single pulse laser to reduce heat damage. Focusing the laser directly on target cells completely destroyed the cells with large traces of burned remnants ( figure S1). To weaken the interaction between the laser pulse and the cell, we adjusted the focus of the laser to the inside of the glass slide and deliver out-focused pulses to the target cell ( figure S1). To determine the optimal out-focusing depth, we tested multiple focuses from surface (0um) to 250um away from the surface into the depth of the glass slide. We heuristically found that 40um out-focusing leads to localized splintering of the glass surface and isolation of targeted cells with minimum damage (figure S1).

For ITO glass experiments, infrared laser was exposed to the target area, evaporating the ITO layer and discharging the target cells of the region. 8 strip PCR tube caps were used as cell retrievers and were pre-treated with air plasma for 30 seconds to make the surface hydrophilic. We mixed 0.5 uL of proteinase K to 7ul of Sample Buffer provided in the genome amplification kit. This mixture was preloaded on the plasma treated retrieving cap. After isolating cells, the cap was connected to a PCR tube and spin down for 1 minute on a mini centrifuge. Enzymatic lysis was then performed on a 50°C heated thermo-cycler for 1 hour. Proteinase K was then inactivated by incubating at 70°C for 10 minutes.

### Whole genome amplification and next generation sequencing

Lysed samples were then incubated at 95°C for 3 minutes to denature double strand DNA and immediately cooled to 4°C to anneal random hexamers. Samples were then mixed with a pre-mixture of 1uL of denaturation solution plus 1uL of neutralization buffer, 9uL of reaction buffer, 1uL of MDA enzyme mix, and 0.2ul of SYBR green I (diluted 1000 times in nuclease free water). Samples were then incubated in a qPCR cycler at 30°C for 90 minutes for MDA reaction, heated to 65°C for 10 minutes to inactivate the enzyme, and cooled to 4°C. The amplified products were purified with AMPure XP kit as suggested by the producer. Amplification quality of each sample was roughly measured by performing PCR validation against 8 selected SNPs across the genome. Among the 8 SNPs, 6 were homologous SNP sites for HL60, 1 was a heterologous SNP for HL60, and 1 was a reference specific site. Validity of the primer design was verified by performing PCR against bulk genomic DNA purified from 10^6^ HL60 cells. We selected samples that produced successful PCR amplicons from all 8 SNP sites for downstream NGS analysis. NGS library was prepared with a SPARK DNA Sample Prep kit and sequenced by a local service company (Celemics, Inc., Korea) using Illumina Miseq machine (with a 250bp paired end reading mode).

### Leukemia panel

Leukemia target panel was designed by prof. Lee at SNUH. The panel contains 643 SNPs where 205 are HL60 specific, 171 are K562 specific, and 267 target both cell lines. From this original panel, we generated a refined panel by collecting SNPs that existed in our cultured cell line samples using bulk genomic DNA target sequencing.

### Genome coverage and CNV analysis

Genome coverage was plotted by normalized sequencing depth. We used a fixed bin size of 9000 bp, resulting in 343,964 bins along the human whole genome (hg18). From a group of randomly selected reads, each read was assigned to its corresponding genome bin. The number of bins containing more than one aligned read was normalized by the total number of bins to calculate the normalized coverage. CNV was computed based on a public method^25^. We neglected the Y chromosome aligned reads.

### Storage condition test

Three blood smear replicates were prepared from a Down syndrome patient’s blood sample and a CATCH 22 patient’s blood sample. Each slide was treated with a different storage condition before cell isolation and WGA experiments. For each sample, one smear was intentionally put under 20 cycles of −20°C freezing and thawing. The other two smears were either stored at room temperature for 8 days or −20°C for 9 days. From each smear we isolated 5 cells, 25 cells per sample with 3 replicates and 100 cells per sample with two replicates. After WGA, we performed 8 selected loci PCR validation to crudely measure WGA quality of each sample.

## Results

### Laser isolation of cells on conventional glass slide

Most researchers and pathologists observe thin slices of tissues or single cell layers of blood samples attached on conventional slide glasses. Therefore, we first aimed to isolate targeted cells on bare glass slides by selectively ablating the glass segment carrying the targeted cell(s). We utilized our previously developed custom LMD device that generates shape-controlled infrared laser pulses^26,27^. In contrast to UV based laser microdissection, our infrared based method causes no damage to the nucleic acids. The shape and size of the optic front of the pulse is controlled by adjustable optical field stops.

However, the isolated cells were damaged by the high power of the nanosecond laser **(Figure S1)**. This is because nanosecond time scale is long enough to occur extra radiation, ionization, vaporization and convection which decreases absorbance constant of the glass^28^.

### Laser isolation of cells on ITO glass slide

Next, we devised a glass slide coated with a sacrificial layer made of indium tin oxide (ITO) **(Figure 1).** This sacrificial layer, which is conventionally used as electrical path patterning, has a high energy absorption rate and thus prevents cell damage. It is also transparent and sustains clear visualization of the samples. Also, compared to commercial LMD products that utilizes polyethylene naphthalene (PEN) films, ITO glasses do not suffer from buckling, tearing, or cause sample visibility disturbance. Importantly, ITO has higher energy absorbance rate of IR light than visible light^29^. And since nucleic acids have lower absorbance at visible- or infrared-light compared to UV^30^, isolating cells on ITO glass slides with IR laser would be ideal.

**Figure 1.**
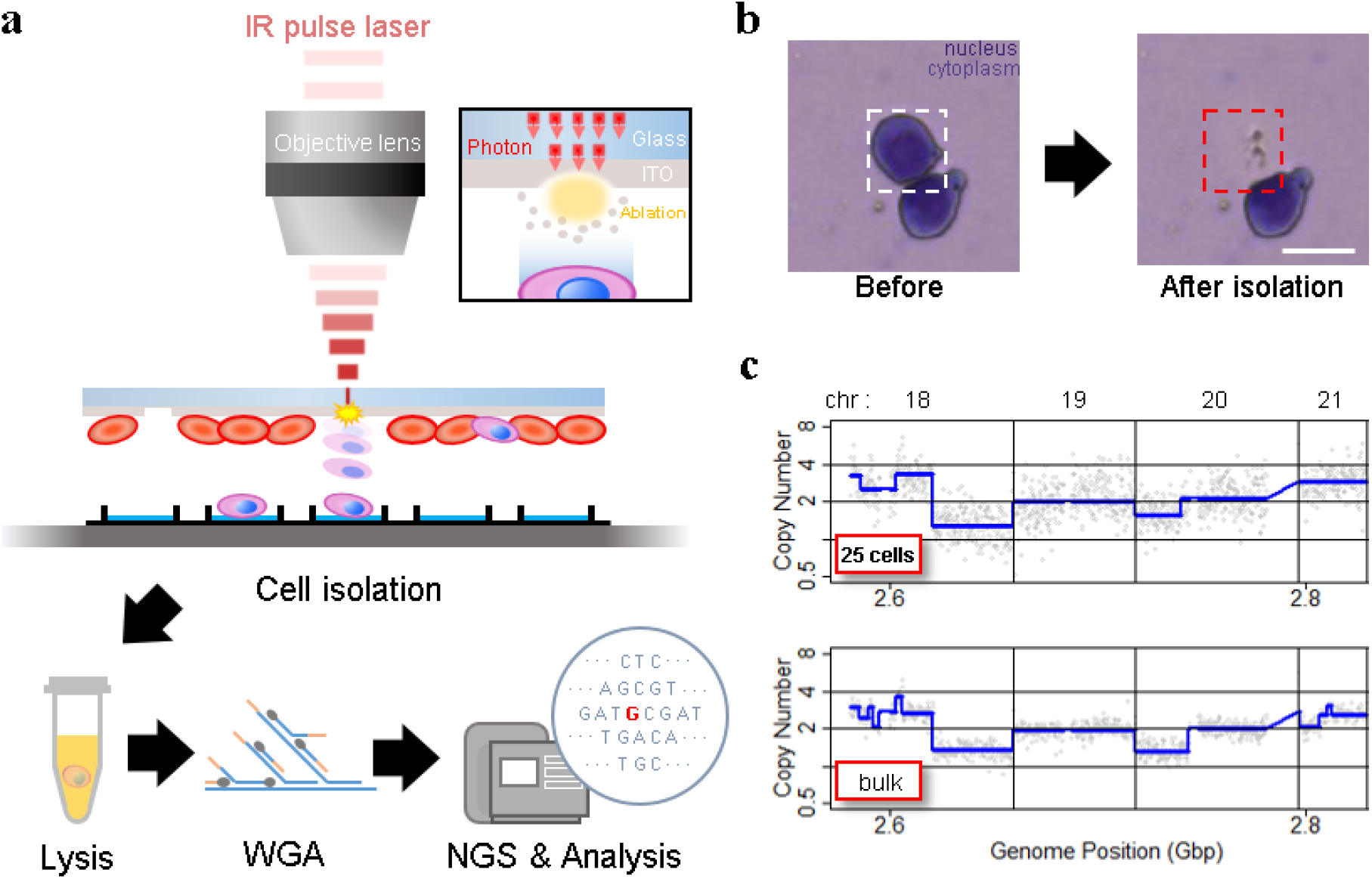
Experiment set-up and representative results. a. Schematic of the IR pulse-laser based cell isolation system and the downstream workflow. b. Representative result of isolating a single cell from a Giemsa stained HL60 cell line sample. c. Representative plot of copy number analysis of K562 cells. The copy number profile obtained from 25 laser-isolated cell sample is comparable to the profile from bulk genomic DNA.

**Figure 2.**
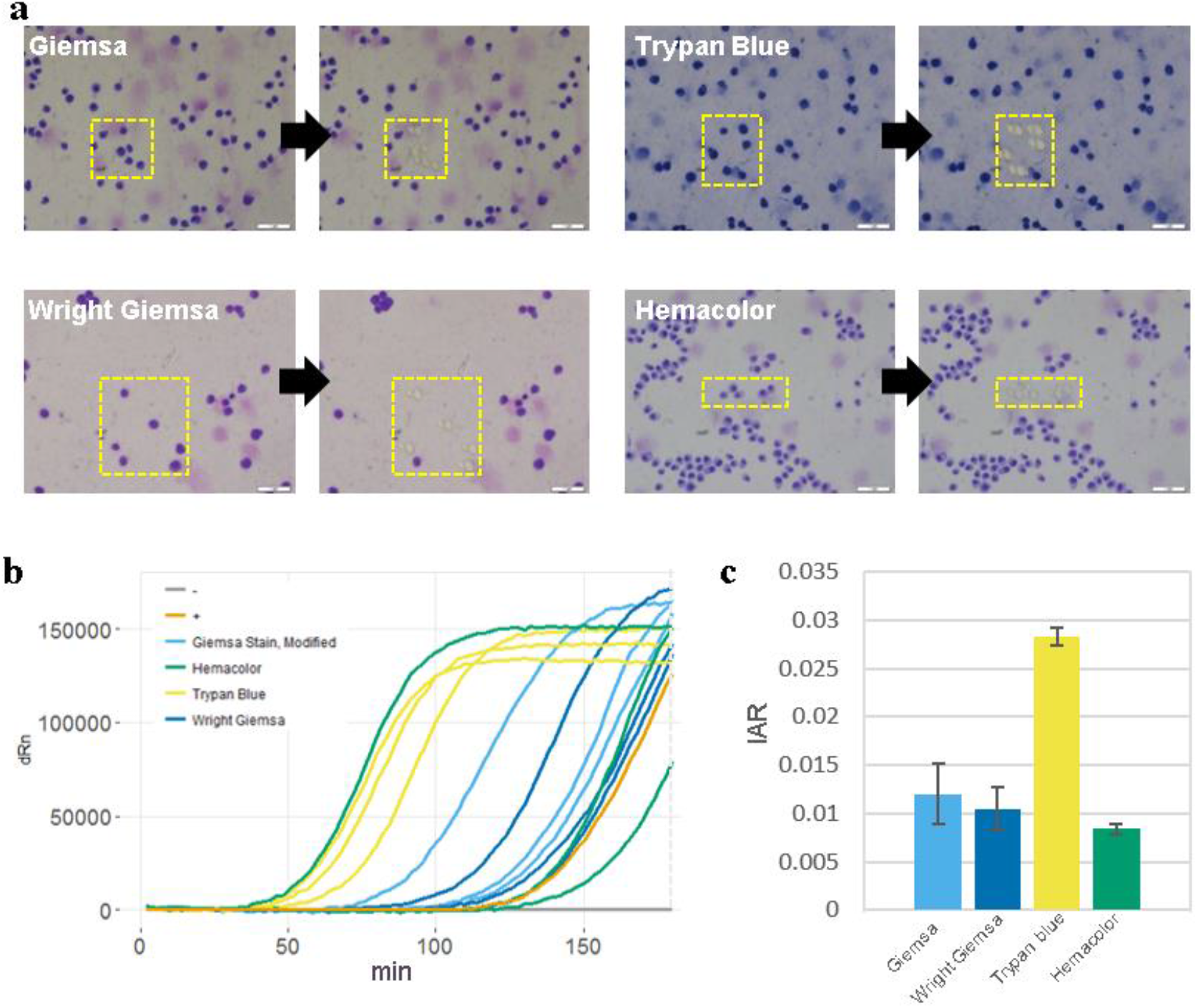
Cell staining protocol affects whole genome amplification efficiency. **a.** Images of cells stained with different staining protocols and smeared on ITO glasses. Yellow boxes indicate laser-targeted cells before (left) and after (right) laser isolation. Scale bars are 50um. **b.** qPCR plot of the whole genome amplification of 5 cell samples. Positive control (+) is a 6pg of purified genomic DNA from bulk HL60 cells. Negative control (-) is a non-template control. **c.** Calculated initial amplification rates (IARs) from qPCR plots in (b).

To test this new approach, we prepared HL60 cell line smears on ITO glasses and stained with Giemsa, which is a widely used hematological staining material. For both ITO layer thickness of 150nm and 300nm, laser isolated cells were retrieved without significant damage **(figure S2)**, suggesting that the scheme works without critical dependency on ITO thickness. In case where targeted cells are not well separated from background cells, decreasing the laser spot size while increasing laser power was sufficient to ensure isolation of target cells only. This indicates that our ITO glass based LMD provides superior target specificity over conventional LMD.

To verify that laser isolated cells are safely retrieved to retrieving cargos (such as slide glass or tube cap), we decided to analyze the dispersity of isolated cells. For this, we prepared H&E stained HL60 cells smeared on an ITO glass and then retrieved one hundred laser isolated cells onto a glass slide. The retrieved cells formed a cluster on the glass slide that was placed 1mm away from the smear sample slide. Dispersity was calculated by measuring the average distance between each retrieved cell and the center (mean or median) of the cluster. Out of the 100 isolated cells, 91 were retrieved on the glass slide. Of those cells, 89 cells were within 1mm radius (or 6σ of a normal distribution). 99% of observed cells were found within 1.25mm radius **(figure S3)**. Since a typical 96 well or PCR tube caps have a diameter around 9mm, our device ensures robust cell isolation into retrieving cargos.

### Genome amplification of retrieved cells

High quality whole genome amplification (WGA) is essential when dealing with low genome input. We modified the lysis and WGA protocol of a commercial kit (GE’s Illustra Genomiphi V2 DNA amplification kit) to increase amplification efficiency. We chose WGA uniformity (i.e. hexamers for MDA annealing uniformly across the whole genome) as an important criteria of high quality WGA. So, with a fixed genomic DNA input, we assumed that more uniform WGA would lead to higher initial amplification rate (IAR, the number of elongating amplicons) measured by real time PCR **(figure S4)**. Amplification rate during subsequent exponential phase should be indistinguishable between high IAR samples and low IAR samples because the intrinsic strand displacement and renewed hexamers annealing characteristic of MDA. All together, we predicted that when measured by real-time PCR, higher amplification uniformity would lead to higher IAR, which would lead to faster entrance to the exponential amplification phase^31^.

To realize this increase in low cell input WGA efficiency, we first replaced alkaline cell lysis with proteinase K based lysis method. Although alkaline lysis method is widely applied to blood cell analysis, in our experience, the WGA efficiency of low DNA input samples (10 cells per sample) were inconsistent among replicates **(figure S5).** We suspected that alkaline lysing may have fragmentized long genomic DNA strands and hinder uniform amplification^32^. We found that by using proteinase K based cell lysis and increasing the incubation temperature from the recommended 37°C to 50°C, higher WGA efficiency was achievable **(figure S6)**. Adding a tapping or vortex step during lysis further improved WGA efficiency, probably by assisting mechanical shearing of the cell and chromatin components **(figure S6)**. Before the MDA step, we also added a DNA denaturing step by incubating the lysate at 95°C for 3 minutes and then cooling to 4°C^33^. This added denaturing step increased random hexamers annealing efficiency which led to higher WGA efficiency **(figure S7. a)**.

We then applied our customized WGA protocol to HL60 single cells isolated with our laser system. For comparison, we performed WGA with 6pg of purified genomic DNA, which is known to be the equivalent amount of DNA from a single cell. The WGA efficiency was comparable between single cells and purified DNA **(figure S7. b)**. We also performed WGA on single cells retrieved by mouth pipetting and also obtained also similar amplification efficiency **(figure S7. b)**. These results proved that our laser-based isolation doesn’t suffer from partial cell retrieval or loss of genomic DNA.

### Analyzing the effect of cell staining on WGA

Pathological blood cell tests usually require nucleic or cell membrane staining to distinguish different cell types. We applied our WGA protocol to HL60 cell line samples pre-treated with one of four popular cell staining methods to analyze the effects of staining on amplification efficiency.

Unfortunately we were not able to produce high quality DNA amplification from single cells. We suspect that DNA staining molecules might have blocked hexamer annealing and/or polymerase elongation. So instead, we isolated 5 cells per sample and compared the amplification efficiency for 4 different staining methods. We used two criteria; i) the IAR measured from the real-time PCR curve, and ii) percentage of successful target PCR (using WGA products as PCR input). For the second criteria, we prepared a small target PCR panel containing 8 genomic sites across the genome **(table S1).**

The result showed that cell staining generally decreases amplification efficiency **(Figure 3)**. Among the four staining methods, Trypan blue showed the highest IAR and highest PCR success rate, indicating high amplification quality **(figure S8)**. This was expected since Trypan blue stains cell membrane proteins and not the nuclear genomic DNA. However, it is a less popular staining method because cell type recognition usually requires nuclear staining. So although Giemsa, a nuclear staining method, staining showed lower amplification efficiency than Trypan blue, it has an advantage over Trypan blue. Hemacolor, which is known to be a fast and user-friendly staining method, showed the lowest amplification efficiency. Consequently, we decided to use Giemsa staining for further analysis.

**Figure 3.**
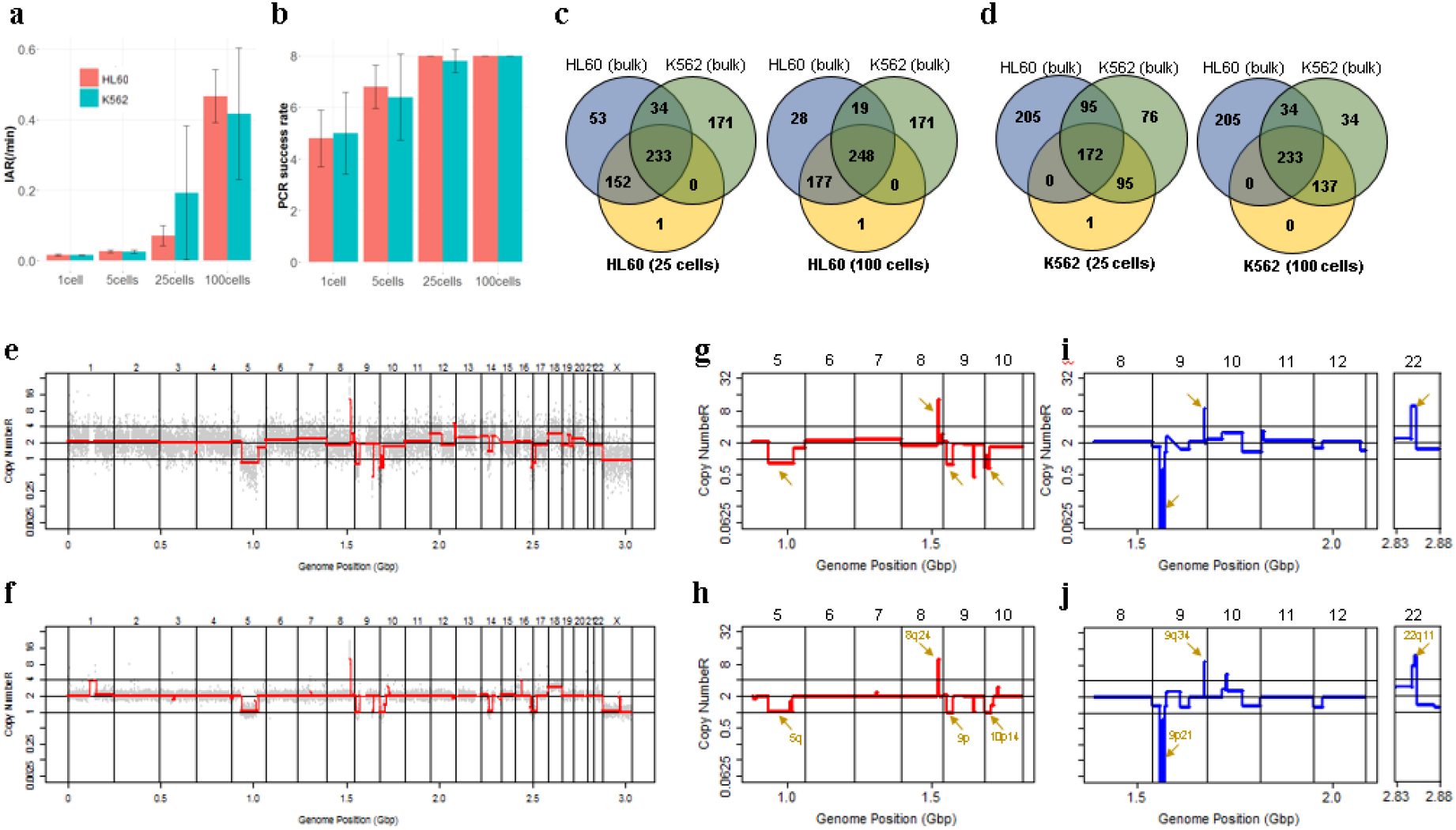
Assessment of WGA quality for various genomic input quantity. 25 cell is sufficient to capture genome-wide profile. **a, b.** Initial amplification rates (a) and PCR success rate (b) from different samples containing 1,5,25,100 cells (n=3). **c,d.** Venn diagrams of the number of detected leukemia panel SNPs. Notice that the 25, 100 cell samples selectively detected their cell-type-specific SNPs regardless of the number of cells per sample. **e,f.** whole genome CN profile obtained from a 25 HL60 cell sample (e) and bulk sample (f). **g-j.** Magnified image of the CN profile from HL60 cells (g,h) and K562 cells (i,j). Notice that the 25 cell profiles (g,i) are highly concordant to the bulk sample profiles (h,j).

### Assessing WGA quality for various genomic input quantity

Since Giemsa staining causes significant sequence information loss during WGA, we decided to assess the minimum number of isolated cells per sample required to retain whole genome coverage. So with WGA products from Giemsa-stained cells, we performed target sequencing and low depth whole genome sequencing to measure allele drop-out rate. In detail, from fixed and Giemsa stained HL60 or K562 cells, we isolated 1, 5, 25, 100 cells per sample, amplified it, and performed our 8 selected loci PCR to crudely measure WGA quality. This PCR panel was targeted at six HL60-specific homozygous SNPs, one heterozygous SNP and one reference allele common for both HL60 and K562. Then for each WGA sample we performed leukemia panel target sequencing (method) and low coverage (0.2x) whole genome sequencing (WGS) to assess genome coverage, and copy number variation (CNV).

As expected, for both HL60 and K562, IAR improved with increasing number of isolated cells per sample **(figure 4. a).** This also showed correlation with panel PCR success rate, where more than 90% success rate was achieved when more than 25 cells were isolated per sample **(figure 4. b)**. Sequencing results of these PCR products also showed that samples with 25 or more cells ensure >90% of true positive SNP call rate **(figure S9)**.

**Figure 4.**
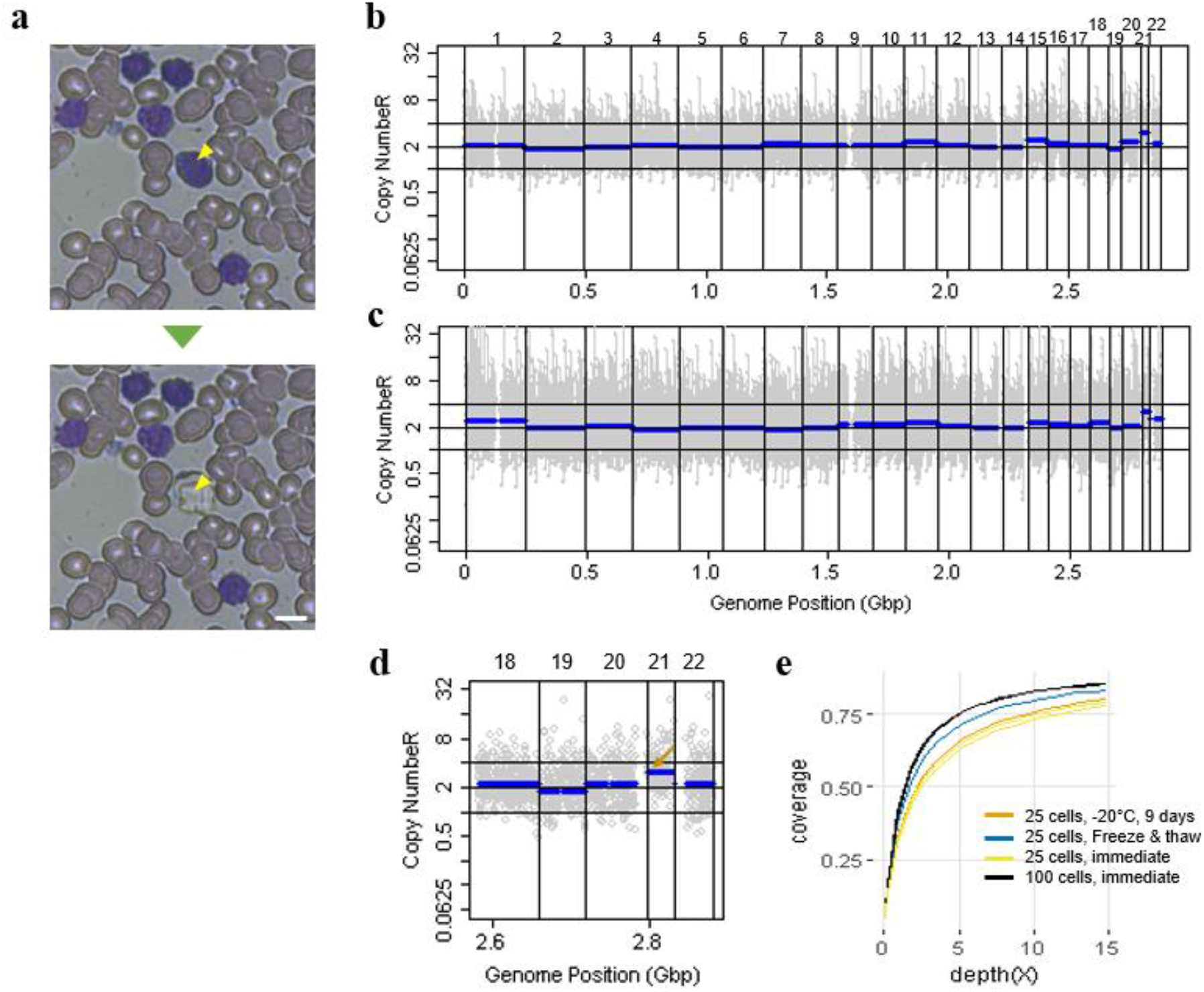
Correct copy number profiling with 25 cells isolated from Down syndrome patient blood smear. **a.** Representative image of single cell isolation from peripheral blood smear. **b, c.** whole genome copy number profile from 100 cell sample (b) and 25 cell sample(c). Cells were laser-isolated immediately after blood smear has been delivered from the hospital. **d.** copy number profile of a 25 cell sample obtained from a freeze-thaw cycled blood smear. The arrow indicates the expected chromosome 21 trisomy. **e.** Genome coverage plot of cells isolated from differently treated blood smear slides. Blood was collected from a Down syndrome patient.

Leukemia panel target sequencing also showed concurrent results. 100 cell samples showed average 87% and 71% true positive rate (TPR) for HL60 and K562 each. This value tended to decline as the number of cells isolated per sample decreased **(figure 4. c, d, S10)**. 5 cell samples showed the lowest TPR. False positive rate, designated here as incident of calling the opposite cell type specific SNP, was non-existent for all experiments **(table S2)**.

With our whole genome sequencing results, we first analyzed genome coverage. Genome coverage of unamplified bulk genomic DNA saturated slightly below 90% whereas the value continuously declined as the number of cells per sample decreased. 25 cell and 5 cell samples showed maximum of 70% and 50% genome coverage **(figure S11. a, b)**. For both HL60 and K562, CNV profiles showed high correlation between bulk sample and 25 cell samples **(figure 4. e, f, figure S11 c, d)**. So even with only 25 cells, characteristic CNVs were detectable such as 5q deletion, 8q24 amplification (related to c-Myc proto oncogene), 9p deletion, and 10p14p11 deletion for HL60 HL60 samples and 9p21 deletion, 9q34 amplification (related to ABL1 proto oncogene), and 22q11 amplification for K562 **(figure 4. g, h)** as reported^34,35^. 5 cell sample CNV profiles showed frequent allele drop outs and prominent amplification bias **(figure S12)**.

In summary, we concluded that isolating more than approximately 25 cells per sample is required to acquire acceptable genome coverage with credible CNV profiling and SNP call of 70~80% TPR. However, higher number of cells per sample or sample pooling may be required for more sensitive mutation or SNP calling experiments.

### Genome scale copy number analysis using pathological blood smears

We then demonstrated CNV analysis using patient blood smears. We collected blood from a Down syndrome (DS) patient and, in less than 12 hours, smeared onto ITO coated glasses and performed fixation and Giemsa staining. We isolated 25 cells, 100cells per sample and performed WGA and low coverage WGS as before. Clearly, the characteristic trisomy of chromosome 21 was observed as expected for both 25cell and 100cell samples (**figure 4. b-d**).

In practice, pathological blood smear slides tend to be stored at room temperature for up to several days before actual pathological observation. This may be acceptable for conventional cell morphology analysis but may hamper genome integrity as well as quality of genomic tests. So for practical purposes, we investigated the effect of blood smear sample storing condition to genome integrity. We tested three conditions; room temperature storage for 8 days, −20°C storage for 9 days, and twenty freeze-thaw cycling. Panel PCR experiment showed that room temperature storage leads to significant reduction in PCR success rate compared to −20°C storage. Repeated freeze-thawing, albeit with 10 minute intervals, seemed to have minimal effect in genome integrity **(figure S13)**. The overall genome coverage of these blood derived cell was lower than previous cell line samples probably due to impurities in the blood plasma. Noticeably, the genome coverage of cell samples from previously stored at −20°C was comparable to freshly isolated cell samples (**figure 4. a**).

In summary, we successfully analyzed CNV profiles of a patient sample using only a small number of blood smear-derived cells and observed that storing blood smears at −20°C is preferred over conventional room temperature storage.

## Discussion

Although routine cytology is a huge source of clinical samples, molecular analysis on these samples still has much room for improvement. In case of genome analysis, conventional cell staining molecules hamper the genome amplification process, leading to frequent allele drop-out and loss of clinical information. This is why LMD is usually performed on non-stained solid tissue sections, and an adjacent tissue section stained with H&E is used as a guide for positioning target cells for dissection^36^. However, this indirect, mirroring technique is not applicable to blood tissue since it lacks the required spatial context.

We developed a new microdissection method and a whole genome amplification protocol that is effective for hematologically stained cell samples. It enables retrieving single cells without genome loss. We also developed a WGA protocol for such stained cells that enables whole genome CN analysis and SNP detection with a minimum number of 25 collected cells, despite low sequencing depth (0.2X). To our knowledge, this is the smallest number cells achieved for WGS from DNA-stained blood cells. In theory, deep (i.e. 100x depth) sequencing enables DNA mutation detection at the single cell level, which was not possible for previous reports where thousands of cells had to be collected to make WGS possible. However, in a more practical view, further improvement will depend on whole genome amplification quality at the single cell level.

The minimum number of cells required for whole genome sequencing will vary with different cell types and staining method. Refining this number based on cell type and staining method of interest may be needed. Future investigation will include applicability of our cell isolation system and WGA protocol to other tissue types such as bone marrow, FISH, and tissue-stamp samples. (**figure S14**). The presented work will find wide applications for immunohistochemical tissue sections and cytological specimens to make novel molecular marker discoveries for the clinic.

## Availability

Related data and methods will be available upon request to the corresponding author.

## Acknowledgement

This research was supported by Global Research Development Center Program through the National Research Foundation of Korea(NRF) funded by the Ministry of Science and ICT(MSIT) (2015K1A4A3047345)

## Tables

**Supplementary table 1.**
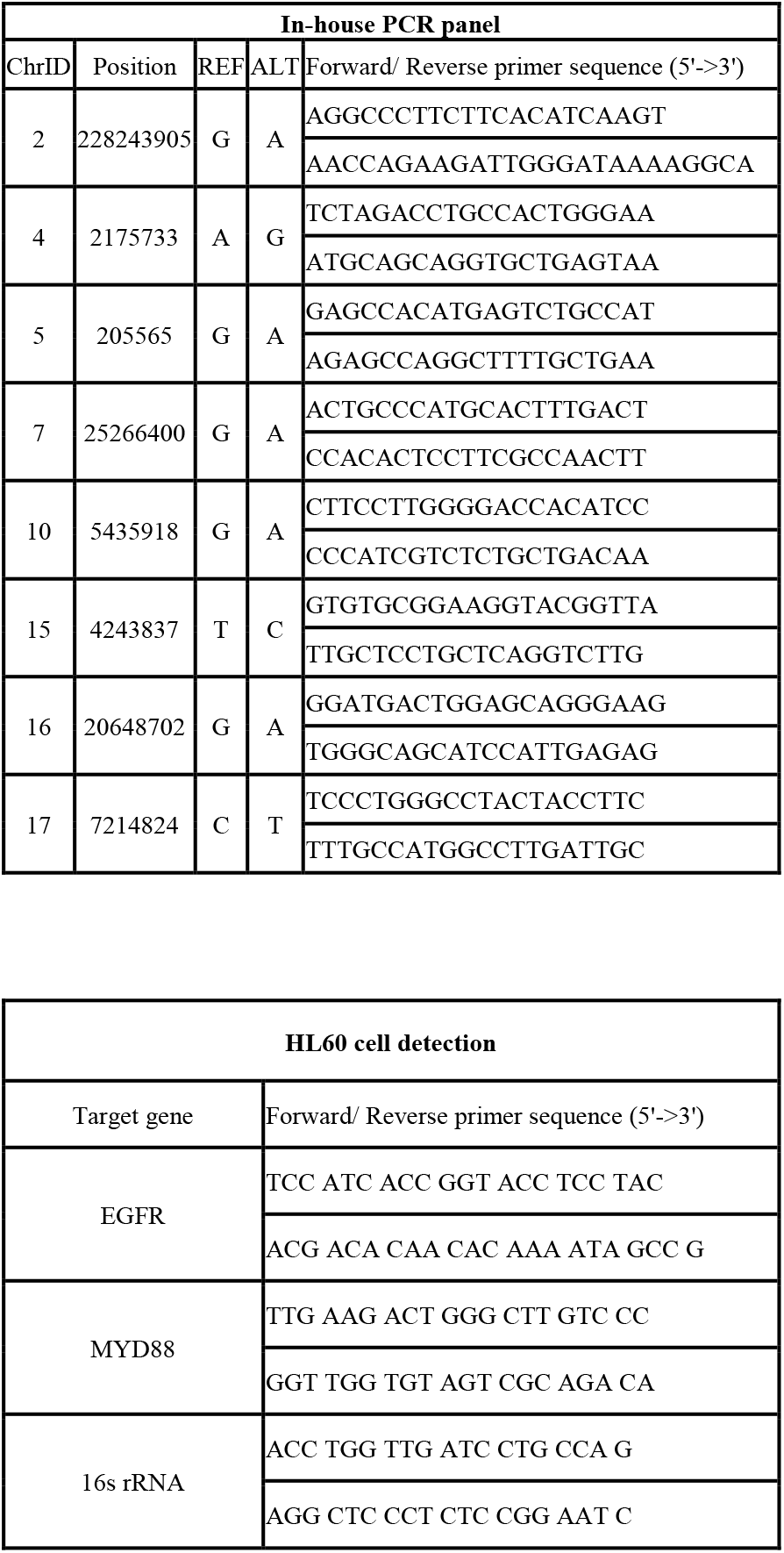
Primer sequence information.

**Supplementary table 2.** Leukemia SNP panel target sequencing result.

|  | HL60 SNPs |  | HL60 specific SNPs |  | K562 specific SNPs |  |  |
| --- | --- | --- | --- | --- | --- | --- | --- |
| Samples | Detected SNPs | % | count | % | Detected SNPs | % | Exception |
| HL60_100-1 | 425 | 90.04% | 177 | 86.34% | 0 | 0.00% | 1 |
| HL60_100-2 | 393 | 83.26% | 157 | 76.59% | 0 | 0.00% | 1 |
| HL60_25-1 | 385 | 81.57% | 152 | 74.15% | 0 | 0.00% | 1 |
| HL60_25-2 | 350 | 74.15% | 137 | 66.83% | 0 | 0.00% | 1 |
| HL60_5-1 | 222 | 47.03% | 73 | 35.61% | 0 | 0.00% | 1 |
| HL60_5-2 | 211 | 44.70% | 74 | 36.10% | 0 | 0.00% | 1 |

|  | K562 SNPs |  | HL60 specific SNPs |  | K562 specific SNPs |  |  |
| --- | --- | --- | --- | --- | --- | --- | --- |
| K562 Samples | Detected SNPs | % | count | % | Detected SNPs | % | Exception |
| K562_100-1 | 370 | 84.47% | 0 | 0.00% | 137 | 80.12% | 0 |
| K562_100-2 | 250 | 57.08% | 0 | 0.00% | 90 | 52.63% | 0 |
| K562_25-1 | 267 | 60.96% | 0 | 0.00% | 95 | 55.56% | 1 |
| K562_25-2 | 375 | 85.62% | 0 | 0.00% | 140 | 81.87% | 1 |
| K562_5-1 | 232 | 52.97% | 0 | 0.00% | 82 | 47.95% | 1 |
| K562_5-2 | 139 | 31.74% | 0 | 0.00% | 41 | 23.98% | 0 |

\* Exception: SNP not detected in bulk control but detected in sample
\* Entire (HL60 and K562) SNP panel: 643
\* HL60 SNPs: 472
\* K562 SNPs: 438
\* HL60 specific SNPs in panel: 205
\* K562 specific SNPs in panel: 171

## Notes

### Competing Interest Statement

The authors have declared no competing interest.

